# *ClassifyITS*: An R Package for assigning taxonomy to fungal ITS sequences using taxon-specific cutoff values

**DOI:** 10.64898/2026.08.31.748297

**Authors:** Quinn S. Moon, Leho Tedersoo, Christian Griebler, Clemens Karwautz, Angela Cukusic, Timothy Y. James

## Abstract

1. Fungi are key drivers of decomposition and nutrient cycling across the globe, yet accurate classification of environmental fungal internal transcribed spacer (ITS) sequences remains challenging. These persistent challenges reflect the variable evolutionary properties of ITS, limited representation of fungal diversity in reference databases, and the application of classifiers originally developed for more conserved prokaryotic markers.
2. Here, we present ClassifyITS, an R package that performs alignment-based taxonomic classification of full length fungal ITS sequences or individual ITS subregions (ITS1 or ITS2) using taxon-specific sequence identity thresholds. In addition to taxonomic assignments, ClassifyITS generates summary statistics and diagnostic visualizations to support interpretation and quality control.
3. Using a deep subsurface fungal ITS dataset containing many poorly characterized taxa, ClassifyITS outperformed the common classifiers SINTAX and DADA2, with higher agreement to expert curated assignments and lower rates of over classifying and under classifying sequences to taxonomic ranks. Across all classifiers and approaches, taxonomic accuracy increased strongly with sequence similarity to the reference database, emphasizing the importance of continued expansion and curation of fungal sequence databases.
4. By providing an accessible and reproducible R based workflow that improves taxonomic classification, ClassifyITS supports more accurate biodiversity monitoring and enhances downstream functional interpretation of fungal communities.

## 1 Introduction

The molecular revolution, particularly the advent of second and third generation sequencing, has transformed our understanding of microbial life. Microorganisms, including fungi, are now known to inhabit environments once considered devoid of life and to drive major nutrient cycling processes across marine, terrestrial, and subsurface ecosystems (Breyer et al., 2025; Coleine et al., 2022; Drake & Ivarsson, 2018). This transformation has been especially important for the study of fungi, which have evolved into an extraordinary diversity of life histories, ranging from mutualistic and saprotrophic to devastating pathogens. Nevertheless, most fungal diversity remains uncultured and is known exclusively from environmental DNA (Hawksworth & Lücking, 2017). Although an estimated 2–10 million fungal species are thought to exist (Hawksworth & Lücking, 2017), only about 5% have been formally described, and approximately 20% of those are represented by sequences in public databases (Abarenkov et al., 2024). Consequently, contemporary fungal surveys rely heavily on environmental DNA to characterize communities and on taxonomic assignments to infer their potential ecological functions (Nguyen et al., 2016; Põlme et al., 2020). Central to this approach is the accurate classification of short marker gene sequences, which underpins inferences about community composition, biogeography, ecological function, and evolutionary diversity. The nuclear ribosomal internal transcribed spacer region, or ITS, is the primary DNA barcode for fungi (Schoch et al., 2012). Despite its widespread use, however, accurate classification of environmental ITS sequences remains challenging, particularly in poorly sampled environments and for deeply divergent or undescribed lineages (Tedersoo et al., 2022).

Both the utility and limitations of ITS as a fungal barcode have been extensively discussed (Kauserud, 2023; Schoch et al., 2012). In practice, ITS sequences are commonly clustered into operational taxonomic units (OTUs), using similarity thresholds between 97% and 99%, or retained as amplicon sequence variants (ASVs) (Nilsson et al., 2019; Tedersoo et al., 2022). Bioinformatics workflows nominate representative ITS sequences that must be assigned taxonomy, which remains one of the most technically limiting steps in community analysis and downstream inference. Several interrelated factors—including the absence of a universal ITS barcode gap, incomplete reference databases, and the limited utility of ITS as a sole marker for phylogenetic inference—complicate fungal taxonomic assignment and must be considered in classifier design (Kauserud, 2023).

Many closely related species within geographically widespread genera, such as *Penicillium* (Lücking et al., 2020) and *Cortinarius* (Garnica et al., 2016), share identical or nearly identical ITS sequences, whereas some fungal lineages exhibit intragenomic ITS variation exceeding 2% (Paloi et al., 2022). This variability underscores the absence of a universal ITS barcode gap across fungi at any taxonomic rank (Kauserud, 2023; Vu et al., 2016, 2019). Recent divergence, incomplete lineage sorting, variation in ribosomal copy number, and demographic history can therefore decouple ITS sequence variation from species boundaries, complicating taxonomic inference at both species and higher taxonomic levels (Kauserud, 2023). These intrinsic limitations are compounded by the incomplete and uneven representation of fungal diversity in public reference databases. Many records are incompletely classified, assigned as *incertae sedis*, or affected by taxonomic or sequence errors (Abarenkov et al., 2024). For example, the UNITE v10.0 all-eukaryote release (Abarenkov et al., 2024) contains 168,030 fungal entries, of which 52,846 are species-level identifications and 34,483 are unique species hypotheses. Relative to estimated global fungal diversity, current ITS databases remain sparse, particularly for recently described or poorly sampled lineages, and often fail to capture the full extent of intraspecific variation within represented species (Hawksworth & Lücking, 2017). Consequently, environmental surveys frequently recover fungal sequences that cannot be classified confidently using existing reference data alone (Guglielmin et al., 2023; Wahl et al., 2018).

The internal transcribed spacer region evolves substantially faster than conserved ribosomal markers such as the 18S and 28S rRNA genes (Nilsson et al., 2019), generally aiding in discrimination among closely related fungal lineages while also resulting in considerable sequence length heterogeneity, insertion and deletion variation, and alignment ambiguity across deeper evolutionary divergences (Kauserud, 2023). Consequently, ITS has limited utility as a sole higher level phylogenetic inference (Orsholm et al., 2026; Tedersoo et al., 2022) and classification frameworks developed conserved prokaryotic markers, including least common ancestor approaches (LCA) commonly applied to 16S rRNA gene datasets, may be poorly matched to its evolutionary dynamics (Barbera et al., 2019; Berger et al., 2011; Y. Wang et al., 2022). Building on a long history of classifier development and improvement (Abarenkov et al., 2018; Gweon et al., 2015; Heeger et al., 2019; Nilsson et al., 2009), accessible, reproducible, and taxon aware approaches can improve the consistency and transparency of fungal ITS classification.

The alignment based Basic Local Alignment Search Tool (BLAST; Camacho et al., 2009) remains valuable for identifying divergent and under sampled lineages because it generates local, gapped alignments that accommodate variation in ITS length and insertion and deletion patterns. The naive Bayesian classifier implemented in DADA2 estimates taxonomic membership from the frequencies of short sequence words in query and reference sequences and uses bootstrap resampling to calculate confidence at successive taxonomic ranks (Callahan et al., 2016; Q. Wang et al., 2007). SINTAX also represents sequences as sets of k mers but assigns taxonomy according to similarity with reference sequences and estimates confidence through repeated subsampling without requiring a trained probabilistic model (Edgar, 2016). In contrast, BLAST reports explicit measures of sequence identity, query coverage, alignment length, gaps, and statistical significance, enabling conservative assessment of distant or ambiguous matches. ClassifyITS extends this alignment-based approach by applying curated, taxon-specific ITS identity thresholds at successive taxonomic ranks and returning the deepest classification supported by both the BLAST evidence and the available reference taxonomy. It thereby standardizes a process that is otherwise commonly performed manually or through laboratory specific workflows while retaining transparent evidence for each assignment. Emerging machine learning and artificial intelligence tools, including MycoAI (Romeijn et al., 2024), offer promising complementary approaches, and ClassifyITS can serve as a reproducible, alignment based method for independently evaluating classifications predicted by these and other classifiers. Although taxon-specific ITS similarity thresholds are increasingly recommended across diverse fungal lineages and habitats (Moon et al., 2026; Tedersoo et al., 2014, 2022), their implementation remains insufficiently standardized, limiting reproducibility, scalability, and accessibility for nonspecialists.

Here, we present ClassifyITS, an R package for fast and reproducible classification of fungal ITS sequences using as input quality controlled BLAST results, taxon informed similarity thresholds (Tedersoo et al., 2021), and consensus based validation. ClassifyITS prioritizes conservative taxonomic assignments and provides user friendly diagnostics that facilitate manual review of ambiguous or highly abundant taxa. The package also quantifies the taxonomic resolution achieved, including the proportions of sequences classified to the genus and species levels, thereby making uncertainty and reference database limitations explicit. Designed as a streamlined workflow for classifying operational taxonomic units from metabarcoding datasets or from individual ITS sequences, ClassifyITS integrates updated, taxon-specific classification rules within a reproducible R framework. It therefore provides a practical tool for fungal ecologists, evolutionary biologists, and microbiome researchers working with ITS sequence data.

## 2 Package Components and Implementation

ClassifyITS is an R package (*R Core Team, 2024)* developed on GitHub and distributed through CRAN. The package requires only input ITS sequences and corresponding BLAST results. The core functions of ClassifyITS can be grouped into three categories: 1) quality control, 2) taxonomic assignment, and 3) summary statistics and visualization. Using the provided BLAST output, the package rapidly performs quality control on both DNA sequences and BLAST results, applies consensus-based taxonomic assignment using customizable taxon-specific cutoff files (Tedersoo et al., 2021), and generates descriptive tables and graphics summarizing the classification run (Fig. 1).

**Figure 1.**
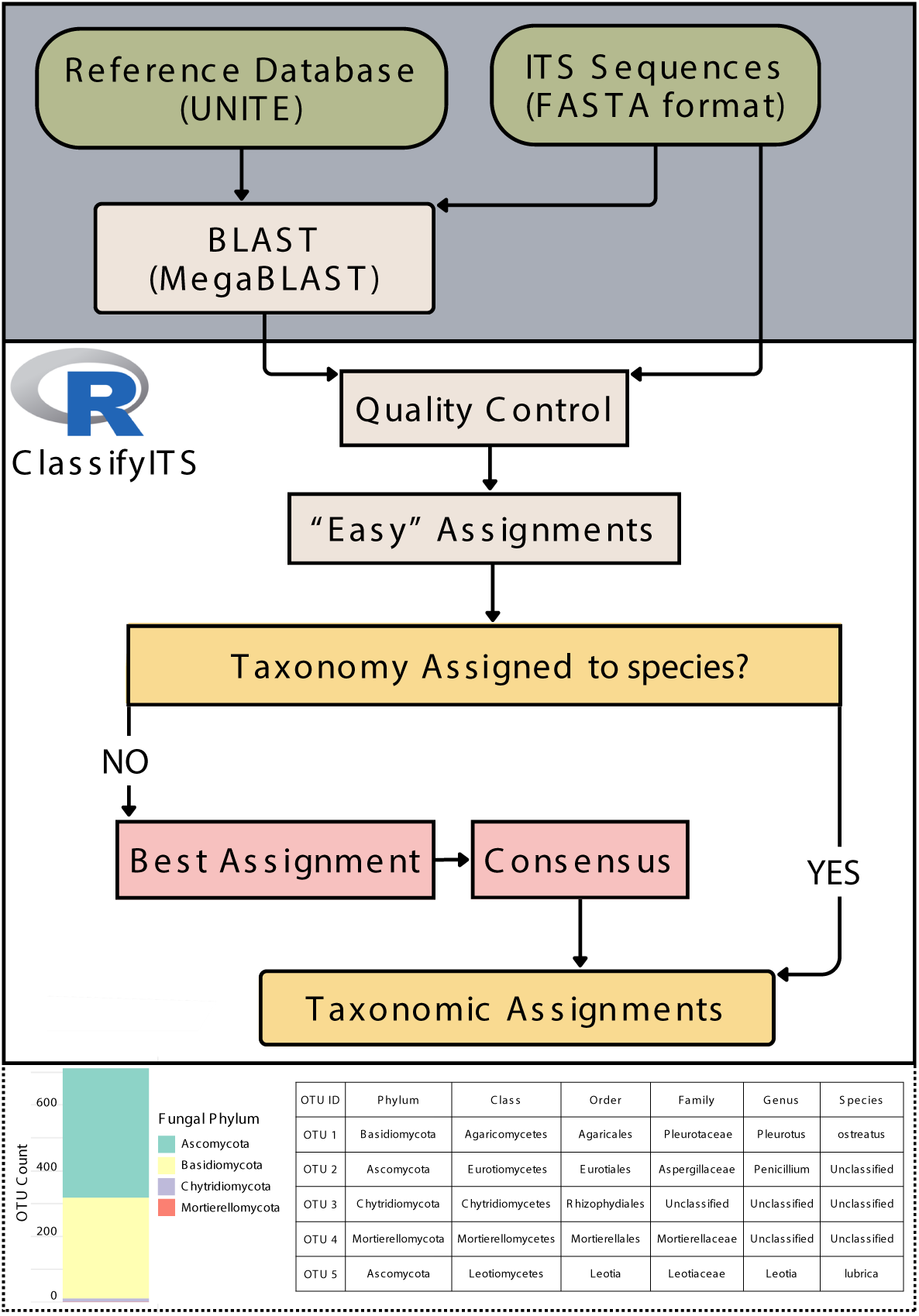
Workflow of the ClassifyITS package. An initial alignment-based BLAST search of ITS sequences against a reference database is performed outside of R, likely on a high-performance computing system, and is shown in the gray box. The resulting BLAST output and corresponding ITS sequences are then passed to ClassifyITS for parsing, quality control, and taxonomic assignment. The pipeline wrapper function ITS_assignment() runs user-provided data through all essential steps of the workflow. When specified, ClassifyITS also generates taxonomic summary outputs, including assignment tables and visualizations. An example taxonomy assignment .csv file and a stacked bar chart summarizing taxonomic composition are shown in the dotted box.

### 2.1 Input data

ClassifyITS is designed to use externally generated BLAST output rather than executing BLAST searches directly within R. Detailed instructions for generating these inputs are provided in the supplementary information and GitHub repository, including step-by-step guidance for both the R interface and the Linux command line. These instructions describe how to download BLAST+, obtain and format the UNITE reference database, and perform the initial BLAST search.

We recommend using the most recent UNITE release, particularly the all-eukaryotes version, because it provides broad taxonomic coverage and includes non-fungal reference sequences. Including non-fungal sequences can improve classification by allowing potential non-target or contaminant sequences to be identified rather than incorrectly assigned to fungi. Nevertheless, reference databases are continually evolving, and future database releases or alternative curated resources, including EUKARYOME (Tedersoo et al., 2024), may also be appropriate depending on the study system and research question.

Within the BLAST+ suite, MegaBLAST is recommended for ClassifyITS workflows because most fungal ITS query sequences are expected to share high nucleotide similarity, typically greater than 95%, with closely related sequences in UNITE. For more divergent nucleotide sequences, such as those commonly observed in Chytridiomycota and Rozellomycota, the computationally slower but more sensitive standard BLASTn task may be appropriate; example compatible parameters are provided in the Supplementary File. The BLAST algorithms, statistical framework, and command line implementation are well documented (Camacho et al., 2009), widely used, and readily deployed on high performance computing (HPC) systems. As BLAST searches can be computationally intensive and are often inefficient to execute within R, ClassifyITS instead focuses on downstream parsing, quality control, and interpretation of externally generated BLAST results.

To support reproducible and high-quality classification, we provide recommended BLAST parameters for use with ClassifyITS, along with instructions for database acquisition and formatting in the Supplementary File. This design allows users to perform computationally demanding similarity searches in an appropriate external environment, such as an HPC cluster, while retaining a standardized and reproducible R based workflow for taxonomic assignment.

ClassifyITS applies taxon-specific sequence similarity thresholds established and continually refined by fungal taxonomic experts (Tedersoo et al., 2021, 2022). These thresholds were developed for full length ITS sequences comprising ITS1, 5.8S, and ITS2 and were calibrated specifically using the UNITE database (Abarenkov et al., 2024), a comprehensive reference resource for fungal ITS sequences. Because UNITE reference records commonly span the complete ITS region, ClassifyITS can also classify sequences representing either ITS1 or ITS2 independently. However, because rates of sequence divergence vary among ITS subregions and taxonomic groups, thresholds derived from full length ITS sequences may be conservative when applied to an individual subregion.

### 2.2 Quality Control

Users provide a representative sequence FASTA file and corresponding BLAST output, which are subjected to quality control using load_and_check(), check_N(), and trim_alignments(). These functions verify required BLAST fields, parse taxonomy strings, remove hits lacking kingdom or phylum level assignments, screen sequences for ambiguous bases, and filter alignments by minimum length. By default, sequences with >1% ambiguous positions are flagged, and alignments shorter than 60% of the median representative-sequence length are removed; both thresholds are user configurable. Records failing quality control criteria are flagged in the final output as these sequences are unlikely to support reliable classification and may be sequencing artifacts (See the Supplementary Information for additional details on contamination checks).

### 2.3 Taxonomic Assignment (Easy, Best, Consensus)

The ClassifyITS classifier implements consensus-based validation to reduce the propagation of erroneous or inconsistent reference annotations into taxonomic assignments. To improve computational efficiency, easy_assignments() first identifies representative sequences that satisfy stringent evidence criteria using quality-controlled BLAST results. Only fungal hits are considered during initial assignment, and hits are ranked by increasing e-value. For sequences with an exact match, the best hit must have 100% sequence identity, family level support must be provided by the two highest ranked hits or a majority of defined hits, and no additional hit with at least 99.5% identity may disagree with the best hit at any taxonomic rank. In the absence of an exact match, the best hit must have at least 98.5% identity, at least two fungal hits must be available, and the best hit must exceed the second best hit by at least 0.5 percentage identity points. If the best and second-best hits agree from kingdom through genus, the taxonomy of the best hit is retained; otherwise, rank specific consensus is evaluated across all fungal hits, with each assignment requiring agreement between the two highest ranked hits or majority support exceeding 50%. Easy assignments are retained only when taxonomy is defined at every rank from kingdom through species and more than half of all BLAST hits support placement within Fungi.

Sequences that do not satisfy the “easy” assignment criteria are processed using best_hit_taxonomy_assignment(), which orders quality-controlled BLAST hits by E value and evaluates each taxonomic rank independently. The highest ranked hit is used by default; however, if its annotation is undefined at a given rank, the corresponding annotation from the second ranked hit may be used when its E value is no more than approximately 1.67 times that of the highest ranked hit. Assignments are evaluated using the most specific applicable taxon informed cutoff, with baseline fungal thresholds applied when no lineage specific rule is available. Kingdom through family are evaluated using E value thresholds, species using percent identity thresholds, and genus using E value, percent identity, or both, depending on the available lineage specific thresholds. Preliminary assignments are subsequently evaluated using consensus_taxonomy_assignment(), which retains a classification at each rank only when it is supported by agreement between the two highest ranked hits or by a greater than 50% majority among defined hits; otherwise, the rank is designated Unclassified. Finally, hierarchical consistency is enforced by designating all ranks below the first Unclassified rank as Unclassified.

### 2.4 Implementation

ClassifyITS includes preloaded taxon-specific cutoffs and default parameter values, allowing users to run the workflow with only the required BLAST output and FASTA file. However, all key parameters can be customized to accommodate dataset specific requirements or alternative cutoff values.

**Table 1.** ClassifyITS parameters for taxonomic assignment. Parameters available for customizing quality control and classification in ClassifyITS are shown; (*) denotes required inputs.

| Parameter | Description |
| --- | --- |
| *blast_file | BLAST results as CSV or TSV, containing top reference hits for each query sequence. |
| *rep_fasta | Representative ITS sequences in FASTA format, used for sequence quality control and alignment length filtering. |
| cutoffs_file | Optional custom CSV file of taxon-specific identity cutoffs; defaults are preloaded. |
| cutoff_fraction | Optional minimum alignment-length threshold, calculated as a fraction of the mean FASTA sequence length. Default = 0.6. |
| n_cutoff | Optional maximum percentage of ambiguous bases allowed per sequence. Default = 1%. |
| outdir | Optional output directory for CSV tables and PDF summary graphics. |
| verbose | Optional setting to print progress and warning messages. |

Detailed installation instructions, example workflows, and quick-start documentation are provided in the supplementary information and on the package GitHub repository. A minimal ClassifyITS workflow requires only a BLAST results file, a representative sequence FASTA file, and, optionally, an output directory:

~~~
’’’
 r
ITS_taxonomy <- ITS_assignment(
 blast_file = “input/blast_results.tsv”,
 rep_fasta = “input/representative_seqs.fasta”,
 outdir = “ClassifyITS_outputs“
)
’’’
~~~

## 3 Additional Features

### 3.1 Flexible taxon-specific cutoffs

As ClassifyITS is designed for application across the fungal kingdom, the default cutoff file provides broad coverage of major fungal lineages. However, taxon-specific thresholds remain unevenly developed, particularly at the species level and among lineages outside Dikarya (Tedersoo et al., 2021; Vu et al., 2016, 2019). To accommodate this uncertainty and incorporate advances in taxonomic knowledge, ClassifyITS supports user defined cutoff files. Users may provide a fully customized cutoff table or modify the default file by adding lineage specific thresholds for taxa of interest. For example, if species level assignment within *Saccharomyces* requires a minimum sequence identity of 99.9%, this threshold can be added to the cutoff file and applied within the ClassifyITS workflow. The required file structure and example implementations are provided in the package GitHub repository and Supplementary Information.

### 3.2 Reproducible Manual Inspection

Alignment-based BLAST searches remain a standard component of fungal taxonomic assessment because these searches provide ranked reference matches, alignment statistics, and database annotations that can be directly evaluated (Lücking et al., 2020; Nilsson et al., 2012). ClassifyITS formalizes this interpretive step by producing reproducible outputs that support targeted manual curation. We recommend prioritizing sequences that remain unclassified at the class level or above, as the recovery of novel species level diversity is common in environmental ITS datasets, whereas unresolved higher-level placement is less frequent and warrants additional scrutiny. High level unclassification for an ITS sequence often reflects reference database incompleteness, particularly for clades known predominantly from environmental DNA and represented in UNITE by incompletely annotated records. However, such patterns may also indicate poor quality sequences, contamination, or potentially novel deeply divergent fungal lineages.

To support transparent manual review, ClassifyITS provides example code for subsetting BLAST results, inspecting ambiguous assignments, and documenting review criteria. This allows users to focus on high priority sequences, such as abundant OTUs, ecologically important taxa, or lineages with uncertain placement, while maintaining a reproducible record of the decision process.

## 4 Validation

To assess the performance of ClassifyITS, we compared its taxonomic assignments with those produced by SINTAX (Edgar, 2016) and the classifier implemented in DADA2 (Callahan et al., 2016), two widely used methods that differ in their implementation and classification strategies. As BLAST-based approaches are expected to be particularly informative for datasets containing taxonomically diverse and undescribed taxa (Tedersoo et al., 2022), we evaluated classifier performance using a fungal ITS2 dataset generated from deep subsurface formation water collected approximately 500 m below the surface and previously shown to contain a substantial novel fungal component (Moon et al., 2026).

### Materials and Methods

Sample collection, library preparation, and OTU generation have been described previously (Moon et al., 2026). Briefly, formation water was collected from deep subsurface wells and filtered through 0.2 µm filters. DNA was extracted using the QIAGEN PowerWater Kit (MoBio Laboratories, Inc., Carlsbad, CA), and fungal communities were characterized through paired end Illumina sequencing of the ITS2 region. Raw reads were merged and denoised using DADA2 (Callahan et al., 2016) and subsequently clustered into OTUs at 98% sequence similarity using VSEARCH (Rognes et al., 2016). The most abundant sequence within each OTU was selected as its representative sequence, resulting in a final validation dataset of 776 OTUs.

For all classification methods, representative sequences were classified against the UNITE all eukaryotes v10.0 database (Abarenkov et al., 2024). Taxonomy was assigned using DADA2 at bootstrap confidence thresholds of 70, 80, and 90, representing the range of thresholds commonly applied in fungal amplicon studies. SINTAX was implemented in VSEARCH using the same confidence thresholds. ClassifyITS was initially run with default parameters to produce a fully automated classification set (ClassifyITS default). A second ClassifyITS classification set was generated following targeted manual inspection of the 41 OTUs that remained Unclassified at the class level or above, using the procedure described above and in the supplementary documentation (ClassifyITS manual).

To establish a reference for evaluating classifier performance, expert curated taxonomic assignments were generated for all representative sequences. For each OTU, fungal taxonomists specializing in zoosporic and aquatic fungi manually examined the top 10 BLAST hits using alignment statistics, taxon-specific identity thresholds (Tedersoo et al., 2021), UNITE annotations, and current knowledge of fungal systematics. These expert curated assignments served as the reference taxonomy for classifier validation. However, taxonomic placement remains inherently uncertain for many environmental ITS sequences, particularly those from under sampled habitats for which reference sequences may be absent, incomplete, or incorrectly annotated, making species level assignments especially provisional. Moreover, this degree of expert curation is impractical for routine datasets, as the manual evaluation of 776 OTUs required more than 150 person hours.

Classifier performance was evaluated using two complementary metrics: classification rate and agreement with the expert curated taxonomy (Fig. 2). Classification rate was calculated as the percentage of OTUs assigned to a given taxonomic rank rather than reported as Unclassified. Agreement was calculated as the percentage of OTUs whose assigned taxonomy matched the expert curated assignment at each rank. As the taxon-specific identity thresholds used by ClassifyITS are currently only developed for fungal classifications below the kingdom level, rank based comparisons below kingdom were restricted to sequences classified as fungi.

**Figure 2.**
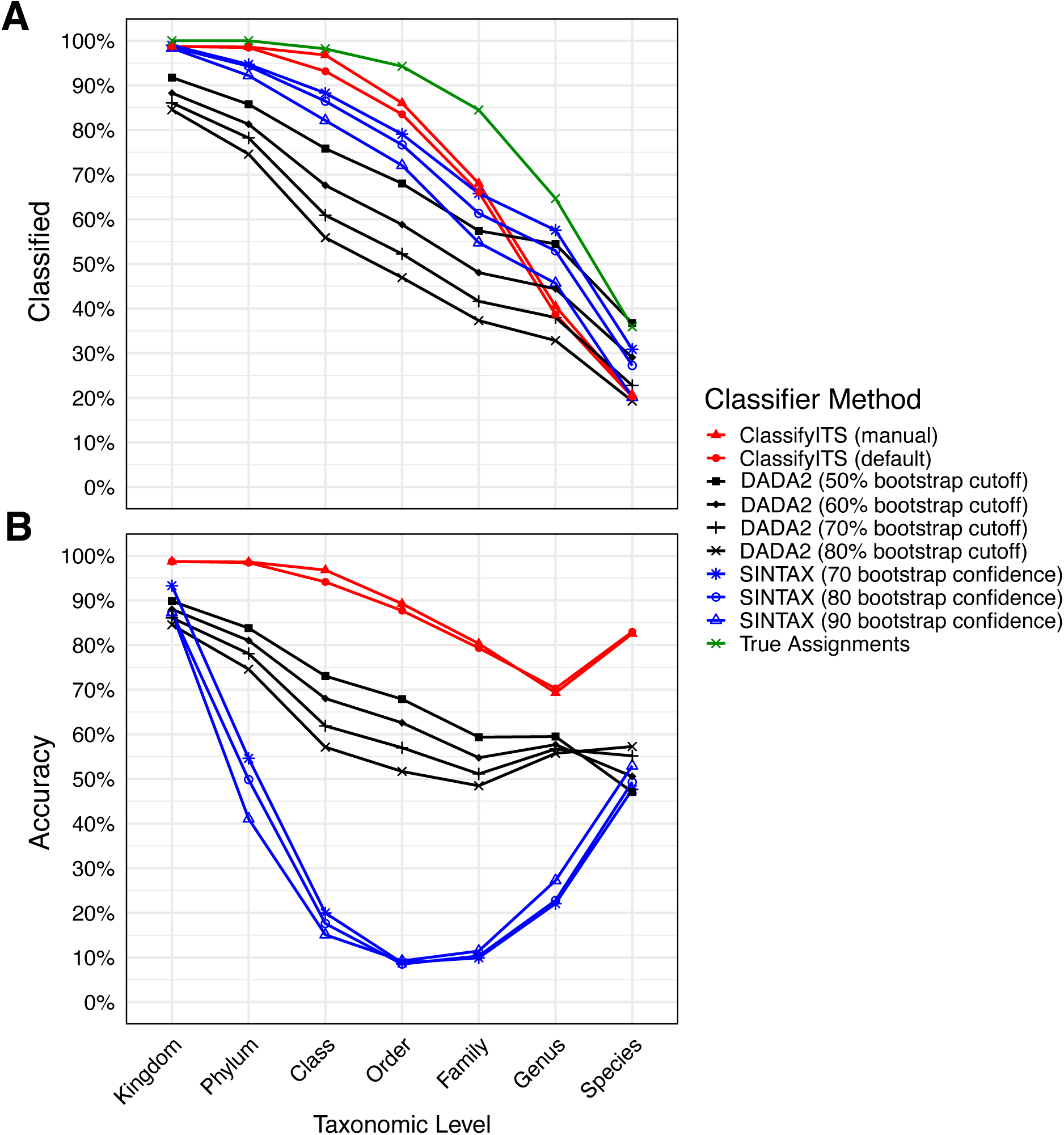
Classification rate and accuracy of ITS taxonomic classifiers across taxonomic ranks. (A) Percentage classified, calculated as the proportion of OTUs assigned a taxonomic label other than Unclassified at each rank. (B) Accuracy, calculated as the proportion of OTU assignments matching the expert curated reference taxonomy. Kingdom level comparisons include all OTUs with reference assignments (n = 734), whereas comparisons from phylum through species include only OTUs assigned to Fungi at the kingdom level (n = 706).

### Results and Discussion

Across all methods, classification rates declined with increasing taxonomic resolution, consistent with the limited representation of environmental fungal diversity in reference databases and the known difficulty of species level ITS assignment (Fig. 2A). ClassifyITS produced relatively high assignment rates at broad taxonomic ranks but became increasingly conservative at lower ranks, resulting in fewer genus and species level classifications than several of the bootstrap-based approaches (Fig. 2B). The pattern of relatively low classification rate in ClassifyITS reflects the design of ClassifyITS which prioritizes conservative, threshold-supported assignments over maximizing species level classifications.

When compared with the expert curated reference taxonomy, ClassifyITS showed the highest overall agreement across all taxonomic ranks (Fig. 2B). The default ClassifyITS workflow closely approximated expert manual inspection, and the manually reviewed ClassifyITS output slightly improved agreement for high level ambiguous OTUs. In contrast, some DADA2 and SINTAX parameterizations produced higher classification rates at lower ranks but lower agreement with expert curated assignments, suggesting a greater tendency toward overclassification in the sampled dataset.

The apparent U shaped pattern in accuracy across taxonomic ranks should be interpreted cautiously. The increase in accuracy at the lowest ranks largely reflects classifiers correctly assigning sequences as Unclassified relative to the expert curated reference, because Unclassified annotations become increasingly common at the genus and species levels. Consequently, conservative methods may exhibit greater agreement with expert assignments at lower ranks by correctly retaining uncertain sequences as Unclassified, despite resolving fewer OTUs to named taxa. Classification rate and accuracy should therefore be considered together when evaluating ITS classifiers.

Overall, these results indicate that ClassifyITS provides taxonomic assignments that are highly concordant with expert curation while requiring substantially less manual effort. The conservative behavior of ClassifyITS is advantageous for datasets containing poorly represented or novel fungal lineages, where forced low rank assignments may introduce systematic error into downstream ecological and evolutionary analyses.

To further characterize classifier performance, we quantified false positive and false negative assignment rates relative to the expert curated reference taxonomy. False positives were defined as cases in which a classifier returned a taxonomic name that differed from the expert curated assignment or assigned a named taxon at a rank designated as Unclassified in the reference. False negatives were defined as cases in which a classifier returned Unclassified at a rank for which the reference contained a named taxonomic assignment. False positive and false negative rates varied substantially among classifiers and parameter settings (Fig. 3). SINTAX produced elevated false positive rates across all tested bootstrap thresholds, with lower thresholds associated with progressively higher false positive rates. These changes in accuracy are consistent with reduced stringency leading to increased assignment of uncertain sequences, particularly in datasets containing divergent or poorly represented taxa. In contrast, DADA2 exhibited comparatively high false negative rates, which increased with higher bootstrap thresholds. This accuracy reflects the conservative behavior of bootstrap-based assignment under stringent confidence criteria, in which uncertain placements are more frequently withheld rather than assigned.

**Figure 3.**
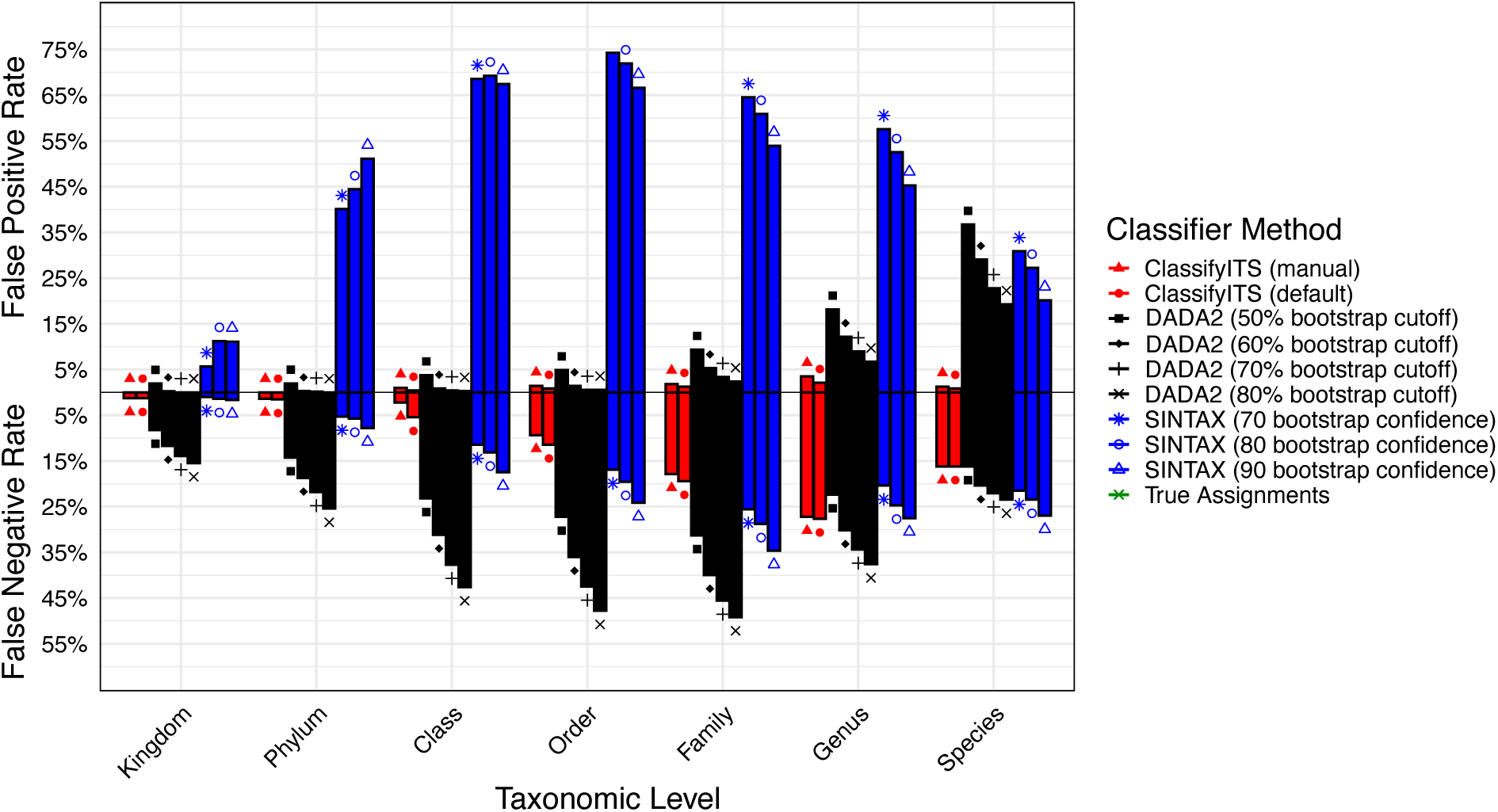
False positive and false negative rates across taxonomic assignment approaches. False positive and false negative rates are shown for each classifier and parameter setting across taxonomic ranks. False positives include both misclassifications and overclassifications relative to the expert curated reference taxonomy, whereas false negatives represent cases in which a classifier returned Unclassified despite a named reference assignment.

ClassifyITS showed lower error rates than all tested SINTAX and DADA2 parameterizations (Fig. 3). Both the default ClassifyITS workflow and the manually inspected ClassifyITS output maintained low false positive (>4%) and false negative rates (>27%) across taxonomic ranks. In particular, false positive rates for ClassifyITS did not exceed 4% at any rank, indicating that the combination of taxon-specific identity thresholds, BLAST-based evidence, and consensus validation effectively limits both misclassification and overclassification while preserving taxonomic resolution.

Assignment accuracy is expected to depend strongly on reference database representation; therefore, we evaluated classifier performance as a function of best-hit similarity to UNITE. For each fungal OTU, the best hit was defined as the highest percent identity BLAST match passing the minimum alignment length filter, at least 60% of the mean representative sequence length, and classified down to at least genus.

Despite the deep subsurface origin of the dataset, 95% of fungal OTUs had a UNITE match greater than 90% identity (Fig. 4A). Across each examined classifier, accuracy increased as best hit identity increased, indicating that regardless of the classifier selected, database proximity is a major predictor of classification success. ClassifyITS performed particularly well for divergent OTUs and reached 100% phylum level agreement when best-hit identity exceeded 85% (Fig. 4B).

**Figure 4.**
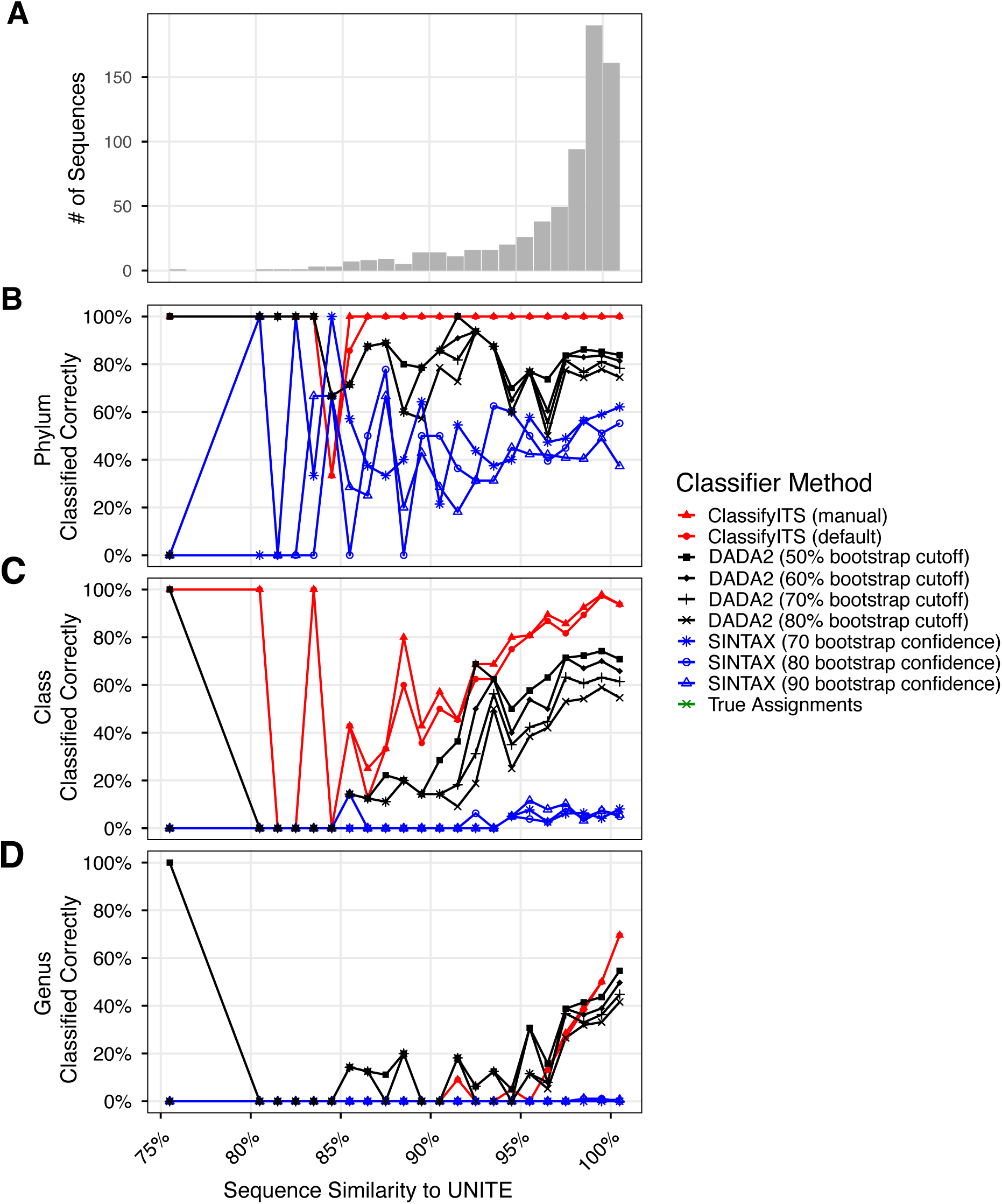
Relationship between reference database similarity and classification accuracy. Percentage of correctly classified fungal OTUs is shown as a function of best-hit percent identity to UNITE. Correct classifications were defined as assignments that were not Unclassified and agreed with the expert-curated reference taxonomy. Analyses included fungal OTUs with BLAST alignments >150 bp and reference assignments resolved to at least class level. Best-hit percent identity was defined as the percent identity of the top BLAST match. In total, 716 fungal OTUs were included; 28 OTUs lacking family-level reference assignments were excluded from family level analyses.

Collectively, the results suggest that ClassifyITS is particularly effective for poorly sampled or divergent fungal lineages and reinforces the importance of continued expansion and curation of ITS reference databases. Since trait databases such as FungalTraits(Põlme et al., 2020) and FUNGuild (Nguyen et al., 2016) rely on family or genus level taxonomy, improved ClassifyITS performance at these ranks may also enhance downstream diversity and functional inference.

## 5 Usage Examples

To further illustrate package use, we developed an example analysis that is included as a package vignette. Incorporating the example workflow as a vignette helps ensure that the code remains compatible with future changes to ClassifyITS and its dependencies.

In addition to the package vignette, we applied the complete ClassifyITS pipeline to the deep subsurface fungal ITS dataset described above and highlight the representative outputs (Fig. 5). The input dataset consisted of 776 representative ITS sequences and 7,760 corresponding BLAST hits. Using default parameters and specifying an output directory, ClassifyITS assigned taxonomy to all sequences in less than 5 seconds on a standard laptop. When an output directory is provided, ClassifyITS creates the directory, if it does not already exist, and writes the primary assignment table, initial_assignment.csv, and summary graphics, combined_taxonomy_graphics.pdf.

**Figure 5.**
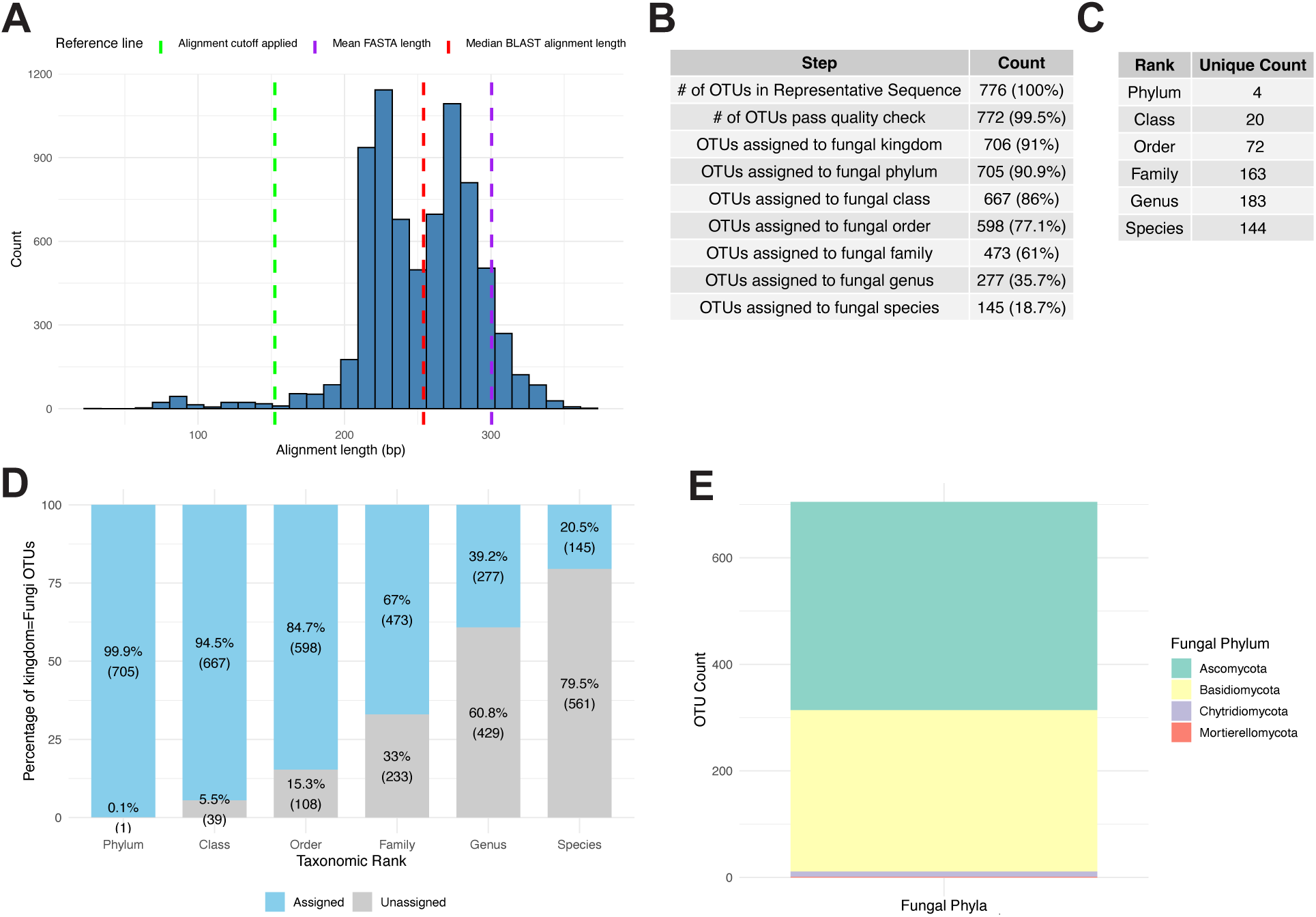
Representative graphical outputs produced by the ClassifyITS pipeline. (A) Distribution of alignment lengths in the user supplied BLAST results. (B) Number and percentage of sequences passing quality control and receiving classifications at major pipeline steps. (C) Counts of unique, defined fungal taxa from phylum through species. (D) Proportion of fungal OTUs classified to a named taxonomic group at each rank. (E) Phylum level taxonomic composition of fungal OTUs. The relative position and order of panels were adjusted for display within a single composite figure.

ClassifyITS returns a tabular output with one row per input sequence and separate columns for the query identifier (qseqid), each taxonomic rank from kingdom through species, and pipeline status (notes). This standardized wide format allows the taxonomy table to be readily joined to an OTU abundance table and imported into downstream community analysis workflows, including phyloseq (McMurdie & Holmes, 2013) and VEGAN (Dixon, 2003). ClassifyITS also provides summary outputs describing run statistics and community composition. In the example dataset, fungal OTUs were dominated by Ascomycota and Basidiomycota, with minor representation from Chytridiomycota and Mortierellomycota.

## 6 Limitations

ClassifyITS provides conservative and reproducible fungal ITS assignments, but its performance remains constrained by the completeness and annotation quality of reference databases, the quality of user-generated BLAST results, and the validity of the underlying taxon-specific thresholds. In particular, the extent to which the thresholds proposed by Tedersoo et al. (2021) accurately delimit taxonomic ranks and can be generalized across the diversity of each fungal lineage remains uncertain and difficult to completely validate. Divergent or poorly represented environmental sequences may therefore remain unresolved even after expert review, underscoring the need for continued database curation and empirical validation and refinement of lineage specific thresholds (Vu et al., 2016, 2019). Finally, ClassifyITS is currently limited to fungi because the implemented cutoff values were developed specifically for fungal ITS sequences; however, appropriate thresholds for other taxonomic groups can be readily added and implemented within the existing framework.

## 7 Conclusion

Despite the rapid expansion of ITS amplicon sequencing for investigating fungal diversity and function, environmental fungal ITS sequences often remain difficult to classify. Accurate characterization of community taxonomic structure can nevertheless provide valuable insight into fungal community function because taxonomy offers an important basis for inferring the potential ecological roles of organisms. Many classification challenges, particularly at the genus and species levels, will persist until reference databases more comprehensively represent fungal genetic diversity and lineage specific barcode gaps are better characterized. ClassifyITS is a user-friendly R package that applies expert derived, taxon-specific thresholds within a reproducible BLAST based workflow. In a validation dataset from a poorly characterized subsurface environment, ClassifyITS outperformed two widely used classifiers, DADA2 and SINTAX, particularly when classifying divergent and undescribed taxa. These results demonstrate that ClassifyITS is a valuable tool for environmental ITS datasets and can be used either as a standalone classifier or alongside complementary approaches to investigate poorly resolved taxa. Improved classification workflows will strengthen efforts not only to document fungal diversity but also to infer the potential ecological functions of microbial communities across Earth’s ecosystems.

## AUTHOR CONTRIBUTIONS

**Quinn S. Moon:** Conceptualization (lead); software (lead); writing—original draft (lead); writing—review and editing (lead). **Leho Tedersoo:** Conceptualization (supporting); writing— review and editing (supporting). **Christian Griebler:** Conceptualization (supporting); writing— review and editing (supporting). **Clemens Karwautz:** Conceptualization (supporting); writing— review and editing (supporting)**, Angela Cukusic:** Conceptualization (supporting); writing— review and editing (supporting), **Timothy James:** Conceptualization (supporting); writing— review and editing (supporting).

## ACKNOWLEDGMENTS

QM acknowledges support from the University of Vienna during the research exchange period.

## DATA AVAILABILITY STATEMENT

The ClassifyITS R package is open source and available via CRAN at (https://cran.r-project.org/web/packages/ClassifyITS/index.html). The development version is hosted on GitHub at (https://github.com/qmoon11/ClassifyITS) and the code used in the validation is available (https://github.com/qmoon11/Validate_ClassifyITS). The static version of the source code and user documentation supporting this article is permanently archived on Zenodo at (10.5281/zenodo.22115679). The sequences used in this study have previously been deposited under NCBI BioProject number PRJNA1346112.

## References

Abarenkov, K., Nilsson, R. H., Larsson, K.-H., Taylor, A. F. S., May, T. W., Frøslev, T. G., Pawlowska, J., Lindahl, B., Põldmaa, K., Truong, C., Vu, D., Hosoya, T., Niskanen, T., Piirmann, T., Ivanov, F., Zirk, A., Peterson, M., Cheeke, T. E., Ishigami, Y., … Kõljalg, U. (2024). The UNITE database for molecular identification and taxonomic communication of fungi and other eukaryotes: Sequences, taxa and classifications reconsidered. Nucleic Acids Research, 52(D1), D791–D797. 10.1093/nar/gkad1039

Abarenkov, K., Somervuo, P., Nilsson, R. H., Kirk, P. M., Huotari, T., Abrego, N., & Ovaskainen, O. (2018). Protax-fungi: A web-based tool for probabilistic taxonomic placement of fungal internal transcribed spacer sequences. 220(2). 10.1111/nph.15301

Barbera, P., Kozlov, A. M., Czech, L., Morel, B., Darriba, D., Flouri, T., & Stamatakis, A. (2019). EPA-ng: Massively Parallel Evolutionary Placement of Genetic Sequences. Systematic Biology, 68(2), 365–369. 10.1093/sysbio/syy054

Berger, S. A., Krompass, D., & Stamatakis, A. (2011). Performance, Accuracy, and Web Server for Evolutionary Placement of Short Sequence Reads under Maximum Likelihood. Systematic Biology, 60(3), 291–302. 10.1093/sysbio/syr010

Breyer, E., Stix, C., Kilker, S., Roller, B. R. K., Panagou, F., Doebke, C., Amano, C., Saavedra, D. E. M., Coll-García, G., Steger-Mähnert, B., Dachs, J., Berrojalbiz, N., Vila-Costa, M., Sobrino, C., Fuentes-Lema, A., Berthiller, F., Polz, M. F., & Baltar, F. (2025). The contribution of pelagic fungi to ocean biomass. Cell, 188(15), 3992–4002.e13. 10.1016/j.cell.2025.05.004

Callahan, B. J., McMurdie, P. J., Rosen, M. J., Han, A. W., Johnson, A. J. A., & Holmes, S. P. (2016). DADA2: High-resolution sample inference from Illumina amplicon data. Nature Methods, 13(7), 581–583. 10.1038/nmeth.3869

Camacho, C., Coulouris, G., Avagyan, V., Ma, N., Papadopoulos, J., Bealer, K., & Madden, T. L. (2009). BLAST+: Architecture and applications. BMC Bioinformatics, 10(1), 421. 10.1186/1471-2105-10-421

Coleine, C., Stajich, J. E., & Selbmann, L. (2022). Fungi are key players in extreme ecosystems. Trends in Ecology & Evolution, 37(6), 517–528. 10.1016/j.tree.2022.02.002

Dixon, P. (2003). VEGAN, A Package of R Functions for Community Ecology. Journal of Vegetation Science, 14(6), 927–930. https://www.jstor.org/stable/3236992

Drake, H., & Ivarsson, M. (2018). The role of anaerobic fungi in fundamental biogeochemical cycles in the deep biosphere. Fungal Biology Reviews, 32(1), 20–25. 10.1016/j.fbr.2017.10.001

Edgar, R. C. (2016). SINTAX: A simple non-Bayesian taxonomy classifier for 16S and ITS sequences (p. 074161). bioRxiv. 10.1101/074161

Garnica, S., Schön, M. E., Abarenkov, K., Riess, K., Liimatainen, K., Niskanen, T., Dima, B., Soop, K., Frøslev, T. G., Jeppesen, T. S., Peintner, U., Kuhnert-Finkernagel, R., Brandrud, T. E., Saar, G., Oertel, B., & Ammirati, J. F. (2016). Determining threshold values for barcoding fungi: Lessons from Cortinarius (Basidiomycota), a highly diverse and widespread ectomycorrhizal genus. FEMS Microbiology Ecology, 92(4), fiw045. 10.1093/femsec/fiw045

Guglielmin, M., Azzaro, M., Buzzini, P., Battistel, D., Roman, M., Ponti, S., Turchetti, B., Sannino, C., Borruso, L., Papale, M., & Lo Giudice, A. (2023). A possible unique ecosystem in the endoglacial hypersaline brines in Antarctica. Scientific Reports, 13(1), 177. 10.1038/s41598-022-27219-2

Gweon, H. S., Oliver, A., Taylor, J., Booth, T., Gibbs, M., Read, D. S., Griffiths, R. I., & Schonrogge, K. (2015). PIPITS: An automated pipeline for analyses of fungal internal transcribed spacer sequences from the Illumina sequencing platform. Methods in Ecology and Evolution, 6(8), 973–980. 10.1111/2041-210X.12399

Hawksworth, D. L., & Lücking, R. (2017). Fungal Diversity Revisited: 2.2 to 3.8 Million Species. Microbiology Spectrum, 5(4), 5.4.10. 10.1128/microbiolspec.FUNK-0052-2016

Heeger, F., Wurzbacher, C., Bourne, E. C., Mazzoni, C. J., & Monaghan, M. T. (2019). Combining the 5.8S and ITS2 to improve classification of fungi. Methods in Ecology and Evolution, 10(10), 1702–1711. 10.1111/2041-210X.13266

Kauserud, H. (2023). ITS alchemy: On the use of ITS as a DNA marker in fungal ecology. Fungal Ecology, 65, 101274. 10.1016/j.funeco.2023.101274

Lücking, R., Aime, M. C., Robbertse, B., Miller, A. N., Ariyawansa, H. A., Aoki, T., Cardinali, G., Crous, P. W., Druzhinina, I. S., Geiser, D. M., Hawksworth, D. L., Hyde, K. D., Irinyi, L., Jeewon, R., Johnston, P. R., Kirk, P. M., Malosso, E., May, T. W., Meyer, W., … Schoch, C. L. (2020). Unambiguous identification of fungi: Where do we stand and how accurate and precise is fungal DNA barcoding? IMA Fungus, 11(1), 14. 10.1186/s43008-020-00033-z

McMurdie, P. J., & Holmes, S. (2013). phyloseq: An R Package for Reproducible Interactive Analysis and Graphics of Microbiome Census Data. PLoS ONE, 8(4), e61217. 10.1371/journal.pone.0061217

Moon, Q. S., Barnhart, E. P., Varonka, M. S., Tomaszewski, E. J., Orozco-Quime, M., Desrosiers, T., Paciorka, I., Carley, M., Schramski, J., Stevenson, B. S., Osburn, M. R., Mohanty, A., Martini, A. M., Stajich, J. E., McIntosh, J. C., & James, T. Y. (2026). Deep subsurface organic-rich shale supports abundant, diverse, and novel fungi. The ISME Journal, wrag184. 10.1093/ismejo/wrag184

Nguyen, N. H., Song, Z., Bates, S. T., Branco, S., Tedersoo, L., Menke, J., Schilling, J. S., & Kennedy, P. G. (2016). FUNGuild: An open annotation tool for parsing fungal community datasets by ecological guild. Fungal Ecology, 20, 241–248. 10.1016/j.funeco.2015.06.006

Nilsson, R. H., Anslan, S., Bahram, M., Wurzbacher, C., Baldrian, P., & Tedersoo, L. (2019). Mycobiome diversity: High-throughput sequencing and identification of fungi. Nature Reviews Microbiology, 17(2), 95–109. 10.1038/s41579-018-0116-y

Nilsson, R. H., Bok, G., Ryberg, M., Kristiansson, E., & Hallenberg, N. (2009). A software pipeline for processing and identification of fungal ITS sequences. Source Code for Biology and Medicine, 4(1), 1. 10.1186/1751-0473-4-1

Nilsson, R. H., Tedersoo, L., Abarenkov, K., Ryberg, M., Kristiansson, E., Hartmann, M., Schoch, C. L., Nylander, J. A. A., Bergsten, J., Porter, T. M., Jumpponen, A., Vaishampayan, P., Ovaskainen, O., Hallenberg, N., Bengtsson-Palme, J., Eriksson, K. M., Larsson, K.-H., Larsson, E., & Kõljalg, U. (2012). Five simple guidelines for establishing basic authenticity and reliability of newly generated fungal ITS sequences. MycoKeys, 4, 37–63. 10.3897/mycokeys.4.3606

Orsholm, J., Zito, A., Somervuo, P., Harrison, J. P., Koskela, M., Ovaskainen, O., Braga, M. P., Chazot, N., Roslin, T., & Furneaux, B. (2026). Discovering the unseen: A performance comparison of taxonomic classification methods for unknown DNA barcodes. Methods in Ecology and Evolution, 2041–210X. 10.1111/2041-210x.70358

Paloi, S., Luangsa-ard, J. J., Mhuantong, W., Stadler, M., & Kobmoo, N. (2022). Intragenomic variation in nuclear ribosomal markers and its implication in species delimitation, identification and barcoding in fungi. Fungal Biology Reviews, 42, 1–33. 10.1016/j.fbr.2022.04.002

Põlme, S., Abarenkov, K., Henrik Nilsson, R., Lindahl, B. D., Clemmensen, K. E., Kauserud, H., Nguyen, N., Kjøller, R., Bates, S. T., Baldrian, P., Frøslev, T. G., Adojaan, K., Vizzini, A., Suija, A., Pfister, D., Baral, H.-O., Järv, H., Madrid, H., Nordén, J., … Tedersoo, L. (2020). FungalTraits: A user-friendly traits database of fungi and fungus-like stramenopiles. Fungal Diversity, 105(1), 1–16. 10.1007/s13225-020-00466-2

R Core Team. R: A language and environment for statistical computing. R Foundation for Statistical Computing, Vienna, Austria. Https://www.R-project.org/ 2024. (n.d.). [Computer software].

Rognes, T., Flouri, T., Nichols, B., Quince, C., & Mahé, F. (2016). VSEARCH: A versatile open source tool for metagenomics. PeerJ, 4, e2584. 10.7717/peerj.2584

Romeijn, L., Bernatavicius, A., & Vu, D. (2024). MycoAI: Fast and accurate taxonomic classification for fungal ITS sequences. Molecular Ecology Resources, 24(8), e14006. 10.1111/1755-0998.14006

Schoch, C. L., Seifert, K. A., Huhndorf, S., Robert, V., Spouge, J. L., Levesque, C. A., Chen, W., Fungal Barcoding Consortium, Fungal Barcoding Consortium Author List, Bolchacova, E., Voigt, K., Crous, P. W., Miller, A. N., Wingfield, M. J., Aime, M. C., An, K.-D., Bai, F.-Y., Barreto, R. W., Begerow, D., … Schindel, D. (2012). Nuclear ribosomal internal transcribed spacer (ITS) region as a universal DNA barcode marker for Fungi. Proceedings of the National Academy of Sciences, 109(16), 6241–6246. 10.1073/pnas.1117018109

Tedersoo, L., Bahram, M., Põlme, S., Kõljalg, U., Yorou, N. S., Wijesundera, R., Ruiz, L. V., Vasco-Palacios, A. M., Thu, P. Q., Suija, A., Smith, M. E., Sharp, C., Saluveer, E., Saitta, A., Rosas, M., Riit, T., Ratkowsky, D., Pritsch, K., Põldmaa, K., … Abarenkov, K. (2014). Global diversity and geography of soil fungi. Science, 346(6213), 1256688. 10.1126/science.1256688

Tedersoo, L., Bahram, M., Zinger, L., Nilsson, R. H., Kennedy, P. G., Yang, T., Anslan, S., & Mikryukov, V. (2022). Best practices in metabarcoding of fungi: From experimental design to results. Molecular Ecology, 31(10), 2769–2795. 10.1111/mec.16460

Tedersoo, L., Hosseyni Moghaddam, M. S., Mikryukov, V., Hakimzadeh, A., Bahram, M., Nilsson, R. H., Yatsiuk, I., Geisen, S., Schwelm, A., Piwosz, K., Prous, M., Sildever, S., Chmolowska, D., Rueckert, S., Skaloud, P., Laas, P., Tines, M., Jung, J.-H., Choi, J. H., … Anslan, S. (2024). EUKARYOME: The rRNA gene reference database for identification of all eukaryotes. Database, 2024, baae043. 10.1093/database/baae043

Tedersoo, L., Mikryukov, V., Anslan, S., Bahram, M., Khalid, A. N., Corrales, A., Agan, A., Vasco-Palacios, A.-M., Saitta, A., Antonelli, A., Rinaldi, A. C., Verbeken, A., Sulistyo, B. P., Tamgnoue, B., Furneaux, B., Ritter, C. D., Nyamukondiwa, C., Sharp, C., Marín, C., … Abarenkov, K. (2021). The Global Soil Mycobiome consortium dataset for boosting fungal diversity research. Fungal Diversity, 111(1), 573–588. 10.1007/s13225-021-00493-7

Vu, D., Groenewald, M., de Vries, M., Gehrmann, T., Stielow, B., Eberhardt, U., Al-Hatmi, A., Groenewald, J. Z., Cardinali, G., Houbraken, J., Boekhout, T., Crous, P. W., Robert, V., & Verkley, G. J. M. (2019). Large-scale generation and analysis of filamentous fungal DNA barcodes boosts coverage for kingdom fungi and reveals thresholds for fungal species and higher taxon delimitation. Studies in Mycology, 92(1), 135–154. 10.1016/j.simyco.2018.05.001

Vu, D., Groenewald, M., Szöke, S., Cardinali, G., Eberhardt, U., Stielow, B., de Vries, M., Verkleij, G. J. M., Crous, P. W., Boekhout, T., & Robert, V. (2016, September 1). DNA barcoding analysis of more than 9 000 yeast isolates contributes to quantitative thresholds for yeast species and genera delimitation [Text]. Westerdijk Fungal Biodiversity Institute. 10.1016/j.simyco.2016.11.007

Wahl, H. E., Raudabaugh, D. B., Bach, E. M., Bone, T. S., Luttenton, M. R., Cichewicz, R. H., & Miller, A. N. (2018). What lies beneath? Fungal diversity at the bottom of Lake Michigan and Lake Superior. Journal of Great Lakes Research, 44(2), 263–270. 10.1016/j.jglr.2018.01.001

Wang, Q., Garrity, G. M., Tiedje, J. M., & Cole, J. R. (2007). Naïve Bayesian Classifier for Rapid Assignment of rRNA Sequences into the New Bacterial Taxonomy. Applied and Environmental Microbiology, 73(16), 5261–5267. 10.1128/AEM.00062-07

Wang, Y., Korneliussen, T. S., Holman, L. E., Manica, A., & Pedersen, M. W. (2022). ngsLCA— A toolkit for fast and flexible lowest common ancestor inference and taxonomic profiling of metagenomic data. Methods in Ecology and Evolution, 13(12), 2699–2708. 10.1111/2041-210X.14006

