## Supplemental File for "*ClassifyITS*: An R Package for assigning taxonomy to fungal ITS sequences using taxon-specific cutoff values"

### Table of Contents

#### Additional Methods Details


1. Data Preparation
2. Install ClassifyITS
3. Quick Start, Run ClassifyITS
4. Inspecting ClassifyITS outputs
5. Screening for human contamination

==================================================================

1. **DATA PREPARATION**

==================================================================

ClassifyITS does not run NCBI BLAST searches or build reference databases. These steps are already well implemented and should be run on HPC (high performance computing cluster). You must generate BLAST results externally and provide them as input. If preferred, equivalent R code using system2() is also provided after the shell workflow. Below is a typical workflow to prepare your inputs (run on HPC):

----------------------------------------------------------------------------

Download BLAST+ (Linux Example)

----------------------------------------------------------------------------

[SHELL SCRIPT — BEGIN]

```
wget https://ftp.ncbi.nlm.nih.gov/blast/executables/blast+/LATEST/ncbi-blast-2.17.0+-x64-linux.tar.gz
tar -xzf ncbi-blast-2.17.0+-x64-linux.tar.gz
export PATH=/your/path/to/ncbi-blast-2.17.0+/bin:$PATH
```

[SHELL SCRIPT — END]

----------------------------------------------------------------------------

Download the Latest UNITE Eukaryotic ITS Database

--------------------------------------------------------------------------------

Download the latest UNITE eukaryotic ITS database from:

https://unite.ut.ee/

ClassifyITS does not download the UNITE database automatically. Users should download the appropriate UNITE FASTA file manually from the UNITE website before running BLAST.

It is recommended to use the latest UNITE eukaryotic ITS database for best results. On the UNITE website, choose the "General FASTA release" that matches your analysis needs. For fungal ITS classification, the eukaryotic ITS release is typically appropriate.

For example, you may download a file such as:

[FILE PATH — BEGIN]

UNITE/euk/sh_general_release_dynamic_s_all_19.02.2025.fasta

[FILE PATH — END]

The exact filename may differ depending on the UNITE release date and selected database type. After downloading, place the FASTA file in your project directory or update the paths in the commands below to point to the downloaded file.

For example, if your downloaded file is located at:

[FILE PATH — BEGIN]

UNITE/euk/sh_general_release_dynamic_s_all_19.02.2025.fasta

[FILE PATH — END]

then use that path in the makeblastdb command below.

--------------------------------------------------------------------------------

Build the BLAST Database

--------------------------------------------------------------------------------

[SHELL SCRIPT — BEGIN]

```
makeblastdb \
  -in UNITE/euk/sh_general_release_dynamic_s_all_19.02.2025.fasta \
  -dbtype nucl \
  -out BLAST/uniteITSEuk_db
```

[SHELL SCRIPT — END]

--------------------------------------------------------------------------------

Run MegaBLAST Search (Example Parameters)

--------------------------------------------------------------------------------

[SHELL SCRIPT — BEGIN]

```
blastn \
  -task megablast \
  -query rep-seqs/dna-sequences.fasta \
  -db BLAST/uniteITSEuk_db \
  -out BLAST/megablast_ITS2.tsv \
  -word_size 28 \
  -reward 1 \
  -penalty -2 \
  -gapopen 0 \
  -gapextend 2 \
  -max_target_seqs 10 \
  -outfmt "6 qseqid sseqid pident length mismatch gapopen qstart qend sstart send evalue bitscore stitle" \
  -num_threads 8
```

[SHELL SCRIPT — END]

This will generate a tab-delimited BLAST output file (for example, BLAST/megablast_ITS2.tsv) that can be used as input for ClassifyITS. It will contain the top 10 hits for each query sequence, along with relevant information such as percent identity, alignment length, e-value, and taxonomic information from the UNITE database.

--------------------------------------------------------------------------------

blastn example (slower)

--------------------------------------------------------------------------------

[SHELL SCRIPT — BEGIN]

```
blastn \
  -task blastn \
  -query rep-seqs/dna-sequences.fasta \
  -db BLAST/uniteITSEuk_db \
  -out BLAST/blastn_ITS2.tsv \
  -word_size 11 \
  -reward 2 \
  -penalty -3 \
  -gapopen 5 \
  -gapextend 2 \
  -max_target_seqs 10 \
  -outfmt "6 qseqid sseqid pident length mismatch gapopen qstart qend sstart send evalue bitscore stitle" \
  -num_threads 8
```

[SHELL SCRIPT — END]

--------------------------------------------------------------------------------

Required BLAST Output Format

--------------------------------------------------------------------------------

Make sure you use:

[SHELL COMMAND — BEGIN]

```
-outfmt "6 qseqid sseqid pident length mismatch gapopen qstart qend sstart send evalue bitscore stitle"
```

[SHELL COMMAND — END]

IMPORTANT: Do not change the column order—this is required for ClassifyITS to work!

==========================================================================

OPTION 2: PREPARE INPUTS FROM R

==========================================================================

If you prefer to run the same preparation steps from R, you can use system2() to call the required Linux/BLAST commands. This does not mean ClassifyITS runs BLAST internally; rather, this is an example of how users can prepare the required BLAST output file themselves from an R session or R script. For large datasets, this should still be run on an HPC compute node, for example through an Rscript submitted with a job scheduler such as SLURM.

------------------------------------------------------------------------------

Prepare BLAST Output for ClassifyITS from R

------------------------------------------------------------------------------

[R SCRIPT — BEGIN]

```r
# Number of CPU threads for BLAST
num_threads <- 8

# Input files
unite_fasta <- "UNITE/euk/sh_general_release_dynamic_s_all_19.02.2025.fasta"
query_fasta <- "rep-seqs/dna-sequences.fasta"

# Output directory and files
blast_dir <- "BLAST"
dir.create(blast_dir, recursive = TRUE, showWarnings = FALSE)

db_prefix <- file.path(blast_dir, "uniteITSEuk_db")
blast_output <- file.path(blast_dir, "megablast_ITS2.tsv")

# If BLAST+ is already available on your PATH, these should be found automatically.
makeblastdb <- Sys.which("makeblastdb")
blastn <- Sys.which("blastn")

# Alternatively, if you downloaded BLAST+ manually, specify the BLAST bin directory:
# blast_bin <- "/your/path/to/ncbi-blast-2.17.0+/bin"
# makeblastdb <- file.path(blast_bin, "makeblastdb")
# blastn <- file.path(blast_bin, "blastn")

# Check that BLAST tools are available
```

```
if (!nzchar(makeblastdb) || !file.exists(makeblastdb)) {
  stop("makeblastdb was not found. Install/load NCBI BLAST+ first.")
}

if (!nzchar(blastn) || !file.exists(blastn)) {
  stop("blastn was not found. Install/load NCBI BLAST+ first.")
}

# Check that input files exist
if (!file.exists(unite_fasta)) {
  stop("UNITE FASTA file was not found: ", unite_fasta)
}

if (!file.exists(query_fasta)) {
  stop("Query FASTA file was not found: ", query_fasta)
}

# Build the BLAST database
system2(
  makeblastdb,
  args = c(
    "-in", unite_fasta,
    "-dbtype", "nucl",
    "-out", db_prefix
  )
)
```

```
# Required BLAST output format for ClassifyITS
blast_outfmt <- paste(
  "6",
  "qseqid",
  "sseqid",
  "pident",
  "length",
  "mismatch",
  "gapopen",
  "qstart",
  "qend",
  "sstart",
  "send",
  "evalue",
  "bitscore",
  "stitle"
)

# Run MegaBLAST
system2(
  blastn,
  args = c(
    "-task", "megablast",
    "-query", query_fasta,
    "-db", db_prefix,
    "-out", blast_output,
    "-word_size", "28",
```

```
    "-reward", "1",
    "-penalty", "-2",
    "-gapopen", "0",
    "-gapextend", "2",
    "-max_target_seqs", "10",
    "-outfmt", blast_outfmt,
    "-num_threads", as.character(num_threads)
  )
)

message("BLAST output written to: ", blast_output)
```

[R SCRIPT — END]

If BLAST+ is not already available on your system, it can be downloaded manually before running the R workflow. For example, on Linux:

[R SCRIPT — BEGIN]

```
dir.create("tools", recursive = TRUE, showWarnings = FALSE)

blast_url <- "https://ftp.ncbi.nlm.nih.gov/blast/executables/blast+/LATEST/ncbi-blast-2.17.0+-x64-linux.tar.gz"
blast_tar <- file.path("tools", basename(blast_url))

download.file(blast_url, blast_tar, mode = "wb")
```

```r
untar(blast_tar, exdir = "tools")

blast_bin <- file.path("tools", "ncbi-blast-2.17.0+", "bin")

Sys.setenv(
  PATH = paste(blast_bin, Sys.getenv("PATH"), sep = .Platform$path.sep)
)

Sys.which("blastn")
Sys.which("makeblastdb")
```

[R SCRIPT — END]

--------------------------------------------------------------------------------

Required BLAST Output Format

--------------------------------------------------------------------------------

Make sure you use exactly:

[SHELL COMMAND — BEGIN]

```
-outfmt "6 qseqid sseqid pident length mismatch gapopen qstart qend sstart send evalue bitscore stitle"
```

[SHELL COMMAND — END]

IMPORTANT: Do not change the column order—this is required for ClassifyITS to work!

**Table 1. Recommended parameters for generating BLAST output compatible with ClassifyITS.** The GitHub repository provides additional guidance and example workflows for producing high-quality BLAST results suitable for input into ClassifyITS.

| **Parameter** | **Description** | **Recommendation** |
|---|---|---|
| word_size | Seed length required to start a match | 28 |
| reward | Score added for each matching nucleotide | 1 |
| penalty | Score subtracted for each mismatched nucleotide | -2 |
| gapopen | One-time cost to start a gap | 0 |
| gapextend | Per-base cost to extend an existing gap | 2 |
| max_target_seqs | Database sequences BLAST reports per query | 10 |

Within BLAST outputs, the principal fields used by ClassifyITS are percent identity (pident) and E-value. Percent identity is reported for each high scoring segment pair (HSP), and is calculated as the proportion of identical nucleotide matches across the aligned region:

$$pident = \frac{\#\ identical\ matches}{alingment\ length}\ x\ 100$$

BLAST also reports an E-value for each alignment. Briefly, a raw alignment score, S, is calculated from the alignment using the specified match, mismatch, and gap penalties. This score is then normalized to a bit score, S′, which is used to estimate the expected number of chance alignments with an equivalent or better score:

$$E = mn2^{-S'}$$

where m is the effective query length, n is the effective database length, and S′ is the bit score. Thus, for each reported alignment, the E-value represents the expected number of database matches that would achieve at least the observed score by chance. Since m and n contribute directly to this calculation, E-values are sensitive to both query length and database size, as well as to the quality of the alignment itself.

==================================================================

### 2. DOWNLOAD AND INSTALL ClassifyITS DIRECTLY FROM CRAN

==================================================================

[R SCRIPT — BEGIN]

```
# Install if you don't have it yet
install.packages("ClassifyITS")

# Load the package
library(ClassifyITS)
```

[R SCRIPT — END]

================================================================

### 3. QUICK START AND RUNNING ClassifyITS

================================================================

Once you have installed the package, and prepared your BLAST results as described in Data Preparation, you can run the assignment pipeline with just a few lines:

[R SCRIPT — BEGIN]

```
ITS_taxonomy <- ITS_assignment(
  blast_file = "input/blast_results.tsv",        # Path to BLAST .TSV file
  rep_fasta = "input/representative_seqs.fasta"  # Path to FASTA file containg representative
sequences used to generate the BLAST results
)
```

[R SCRIPT — END]

By default, ITS_assignment() does not write any files. If you'd like the pipeline to also export the assignments table (CSV) and summary graphics (PDF), supply an output directory:

[R SCRIPT — BEGIN]

```
ITS_taxonomy <- ITS_assignment(
  blast_file = "input/blast_results.tsv",
  rep_fasta  = "input/representative_seqs.fasta",
```

```
  outdir     = "outputs"
)
```

[R SCRIPT — END]

While not necessary for a successful run, the assignment process can easily be customized with additional parameters:

[R SCRIPT — BEGIN]

```
ITS_taxonomy <- ITS_assignment(
  blast_file       = "input/blast_results.tsv",
  rep_fasta        = "input/representative_seqs.fasta",
  cutoffs_file     = NULL,       # Optional: custom taxonomy cutoffs CSV file
  cutoff_fraction  = 0.6,        # Optional: fraction for alignment length QC
  n_cutoff         = 1,          # Optional: percent N cutoff
  outdir           = "outputs",  # Optional: output directory (writes CSV/PDF when provided)
  verbose          = FALSE       # Optional: print progress messages
)
```

[R SCRIPT — END]

==================================================================

4. **MANUAL INSPECTION (UNCLASSIFIED OTUs AT PHYLUM OR CLASS LEVEL)**

================================================================================

Depending on your research question and sample size, it is feasible to subset the BLAST results to only those fungal taxa unclassified at phylum or class, and manually inspect and assign taxonomy:

- A small percentage of taxa (up to approximately 10% in novel habitats) may be unclassified at the phylum or class level. This does not automatically indicate discovery of a new order or phylum.
- It is strongly recommended to manually check the BLAST results for these OTUs:
  - If they have strong BLAST hits to known fungal taxa, they may represent novel taxa not well-represented in the database.
- Occasionally, errors in a database (like UNITE ) (e.g., an entry classified as fungi_sp at the phylum level) will prevent assignment, even if other BLAST results indicate a confident placement (such as Ascomycota).

[R SCRIPT — BEGIN]

```
# Read in BLAST results and assignments
blast <- read.table(
  "input/blast_results.tsv",
  sep = "\t",
  header = TRUE
) #user provided BLAST results

assignments <- read.csv(
  "outputs/initial_assignments.csv",
```

```
  stringsAsFactors = FALSE
) #preliminary taxonomy assignments from ClassifyITS


# Subset assignments to fungal OTUs unclassified at class level
unclassified_class_otus <- assignments[
  assignments$kingdom == "Fungi" &
  (is.na(assignments$class) |
   assignments$class == "" |
   assignments$class == "Unclassified"),
  "qseqid"
]


# subset blast results to these OTUs
blast_sub <- blast[blast$qseqid %in% unclassified_class_otus, ]


# View the BLAST results for these OTUs
View(blast_sub)


# You can also export this subset for manual inspection in Excel
write.table(
  blast_sub,
  "outputs/unclassified_class_blast_results.tsv",
  sep = "\t",
```

```
  row.names = FALSE,
  quote = FALSE
)
```

[R SCRIPT — END]

This is the recommended approach, as ClassifyITS is designed to conservatively assign taxonomy and allow for further inspection at relevant taxonomic levels. Manual curation is especially valuable for ambiguous or novel taxa.

**5 Potential Human Contamination**

Amplicon sequencing can recover contaminant DNA, including human associated fungi such as skin associated Malassezia (Rahimlou et al., 2025), particularly in low biomass samples (Fierer et al., 2025). To support contaminant screening, ClassifyITS includes an optional FASTA file of ITS sequences generated from the holotype specimen of commonly encountered obligate or strongly human associated fungal taxa. Users can BLAST representative sequences against this database to flag potential contaminants for further inspection or removal. This feature is intended as a screening tool rather than an automatic filter, and flagged taxa should be interpreted in the context of sample type, study design, and ecological expectations.

### References

Fierer, N., Leung, P. M., Lappan, R., Eisenhofer, R., Ricci, F., Holland, S. I., Dragone, N., Blackall, L. L., Dong, X., Dorador, C., Ferrari, B. C., Goordial, J., Holmes, S. P., Inagaki, F., Korem, T., Li, S. S., Makhalanyane, T. P., Metcalf, J. L., Nagarajan, N., … Greening, C. (2025). Guidelines for preventing and reporting contamination in low-biomass microbiome studies. *Nature Microbiology*, *10*(7), 1570–1580. https://doi.org/10.1038/s41564-025-02035-2

Rahimlou, S., Amend, A. S., & James, T. Y. (2025). Malassezia in environmental studies is derived from human inputs. *mBio*, *16*(6), e01142-25. https://doi.org/10.1128/mbio.01142-25
